# Copper selects for siderophore-mediated virulence in *Pseudomonas aeruginosa*

**DOI:** 10.1101/2021.09.08.459405

**Authors:** Luke Lear, Elze Hesse, Angus Buckling, Michiel Vos

**Affiliations:** European Centre for Environment and Human Health, University of Exeter Medical School, Penryn, Cornwall, United Kingdom, TR10 9FE; College of Life and Environmental Science, University of Exeter, Penryn, Cornwall, United Kingdom, TR10 9FE

**Keywords:** evolution of virulence, opportunistic pathogen, metal detoxification, pyoverdine, coincidental selection

## Abstract

Iron is essential for almost all bacterial pathogens and consequently it is actively withheld by their hosts. However, the production of extracellular siderophores enables iron sequestration by pathogens, increasing their virulence. Another function of siderophores is extracellular detoxification of non-ferrous metals. Here, we experimentally link the detoxification and virulence roles of siderophores by testing whether the opportunistic pathogen *Pseudomonas aeruginosa* displays greater virulence after exposure to copper. To do this, we incubated *P. aeruginosa* under different environmentally relevant copper regimes for either two or twelve days. Subsequent growth in a copper-free environment removed phenotypic effects, before we quantified pyoverdine production (the primary siderophore produced by *P. aeruginosa*), and virulence using the *Galleria mellonella* infection model. Copper selected for increased pyoverdine production, which was positively correlated with virulence. This effect increased with time, such that populations incubated with high copper for twelve days were the most virulent. Replication of the experiment with a non-pyoverdine producing strain of *P. aeruginosa* demonstrated that pyoverdine production was largely responsible for the change in virulence. Therefore we here show a direct link between metal stress and bacterial virulence, highlighting another dimension of the detrimental effects of metal pollution on human health.

## Introduction

Iron is essential for the growth of almost all bacteria as it serves as a cofactor for many enzymes (1, 2). However, as iron most commonly exists in the insoluble form Fe^3+^, it is of relatively low bioavailability in the majority of environments (1-7). It is therefore essential for bacteria to actively sequester Fe^3+^ from the environment (8, 9), with many bacteria producing iron-chelating siderophore compounds to do this (10). Siderophores aid iron recovery by forming extracellular complexes with Fe^3+^ which are then taken up by the cell and reduced to the bioavailable form Fe^2+^ (11). The extracellular nature of siderophores means they can benefit the group around the producer, with only the producer paying the cost of production. As a result, siderophore production can be a cooperative trait subject to social evolution (12, 13). Therefore as increased siderophore production evolves under iron limitation to benefit the producer and its kin, genotypes with reduced production can also evolve to exploit these producers and gain a fitness advantage (12). Due to the importance of iron for microbial growth, one of the first lines of host defence is to withhold iron from invading pathogens (14). This often involves the production of molecules such as transferrin that bind to iron with very high association constants (6, 14-16). Such nutritional immunity can be overcome by pathogens through the production of siderophores, as their very high iron affinity enables ‘stealing’ iron from the host (16). Siderophores are therefore an important virulence factor in both gram-negative-(17-19) and gram-positive human pathogens (20, 21).

In addition to iron, siderophores are known to bind to a range of potentially toxic metals (22). Crucially, these complexes cannot passively diffuse into the cell (*i.e*. not without active use of transport proteins) and consequently the metal is detoxified extracellularly (23). As a result, siderophore production has been shown to increase in the presence of toxic levels of a wide range of non-ferrous metals (2, 3, 9, 22). As many of the metal ions that can be bound by siderophores are essential in small amounts, siderophore production is adjusted in response to metal bioavailability (24), with either active uptake of metal-siderophore complexes or other metal chelating compounds used to acquire these micronutrients when they are scarce (7, 25). Adjustments in production can be due to both genetic changes and phenotypic plasticity, with genetics dictating the limits of siderophore production and phenotypic plasticity used to regulate production within those limits (26). Although the upper limit of production initially increases with metal toxicity, eventually a threshold is reached as the rising metabolic cost of siderophore production can result in selection for reduced production (8) – analogous to the evolutionary response to iron deficiency (27). This cost is further exacerbated when competitors of the producer benefit from the siderophore (12). Consequently, when there is strong selection for siderophore production in the community, such as when toxic metals are at high concentrations, per capita siderophore production can be reduced both through phenotypic and genotypic change (8, 27, 28). Phenotypic changes in production occur quickly in response to alterations in the environment, including the abundance of metals and population densities, and allow the cost of production to be minimised (23, 28).

Despite strong evidence both for siderophores playing an important role in metal detoxification (3, 8, 22, 24) and for siderophore production being an important determinant for bacterial virulence (16, 29, 30), whether the presence of toxic metals in non-host environments affects virulence remains untested. Here we aim to experimentally link these dual effects of siderophore production, by testing the effects of copper on the virulence of the opportunistic human pathogen *Pseudomonas aeruginosa. P. aeruginosa* is of clinical significance capable of surviving in a range of environments including soil, water and fomites in hospitals (31, 32). In clinical settings it is frequently responsible for severe and fatal infections of patients with cystic fibrosis (CF) (33), burns (34), and immunosuppressive illnesses (35). It is important to note that many CF infections are caused by *P. aeruginosa* acquired from the environment (36), and so an understanding how selection for siderophore production outside of the host could alter virulence is of great importance.

The production of pyoverdine, the main *P. aeruginosa* siderophore, has been shown to rapidly evolve (37). We therefore test virulence after both two days (∼7 generations), when little genotypic change is expected to have occurred, and twelve days (∼40 generations), where genotypic changes in production are expected (12, 38). To allow us to exclude phenotypic changes to production, at each timepoint (two and twelve days) populations are transferred to a common garden environment (copper free media) for 24 hours prior to assays. By quantifying pyoverdine production in every population before the virulence assay, we are able to test whether its production is associated with virulence. We then further test for an association by repeating the experiment using a *P. aeruginosa* strain that cannot produce either pyoverdine or pyochelin (*P. aeruginosa*’s secondary siderophore (39)), and testing whether copper has the same effect on virulence. Moreover, we test if this association is the same across two environmentally relevant copper concentrations that span those found in agricultural soil (40) and in water (41).

## Methods

### Experimental design

*Pseudomonas aeruginosa* PAO1 (42) was grown shaking at 180 rpm for 24 hours at 28°C in glass microcosms containing 6mL of King’s medium B (KB; 10g glycerol, 20g proteose peptone no. 3, 1.5 g K_2_HPO_4_,1.5 g MgSO_4_, per litre). After homogenisation by vortexing, 60µL of undiluted culture was added to 18 microcosms containing KB mixed with sterilised copper sulphate (CuSO_4_; Alfa Aesar, Massachusetts, United States) to final concentrations of either 0.0, 0.1 or 1.0 g/L. Inoculated microcosms were kept static at 28°C for 48 hours before being thoroughly homogenised and 60µL (1% by volume) transferred into fresh media (Fig.1). Microcosms were incubated at 28°C, rather than the more common 37°C, in order for our experiment to be more representative of environmental conditions. Transfers occurred every two days for twelve days. To control for the physiological effects of copper stress, on days two and twelve each microcosm was homogenised and 60µL transferred into a common garden environment (6mL of KB medium without copper). These cultures were grown for 24 hours at 28°C before pyoverdine production was quantified and aliquots were frozen at -80°C in glycerol at a final concentration of 25% for virulence assays and to quantify density. We then repeated this experiment with an isogenic mutant strain of *P. aeruginosa* (PAO1Δ*PvdD*Δ*PchEF* (43)) with the ability to produce pyoverdine, and the secondary siderophore pyochelin, knocked out (a non-pyoverdine producing strain).

**Figure 1.**
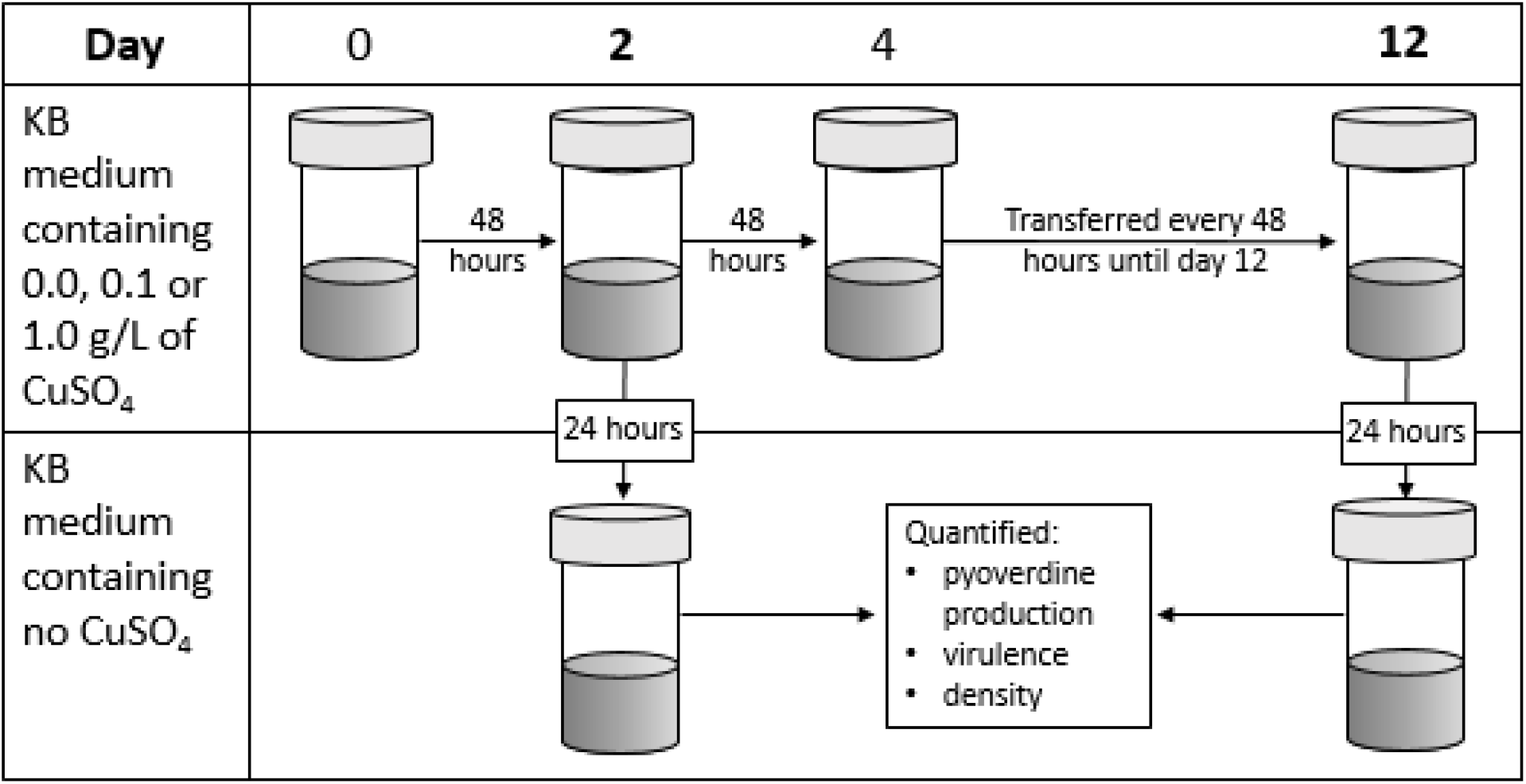
Schematic of the experimental design used to test whether copper selects for siderophore-mediated virulence. Microcosms (n=6 per treatment) containing KB medium at a concentration of either 0.0, 0.1 or 1.0 g/L of copper sulphate (CuSO_4_) were inoculated with *Pseudomonas aeruginosa* (either the pyoverdine producing or non-pyoverdine producing strain), incubated at 28°C and transferred every two days into fresh media. On days two and twelve, cultures were transferred into copper free medium for 24 hours before being homogenised, their per capita pyoverdine production quantified and frozen in glycerol at a final concentration of 25%. Virulence and density assays were performed using defrosted samples.

### Pyoverdine assay

Pyoverdine production was quantified after both two and twelve days. Following 24 hours in a common garden environment, cultures were thoroughly homogenised before 600µL was transferred into a 96 well-plate (200µL into three separate wells). The optical density at 600nm (OD_600_) and the fluorescence at 460nm following excitation at 400 nm of each of the cultures in the 96-well plate was measured using a BioTek Synergy 2 plate reader (BioTek, Vermont, U.S.A.). Pyoverdine, which fluoresces green, is the only culture component quantified using these excitation and emission parameters, with non-producers giving a reading of zero relative fluorescence units (RFU) (4). Each culture was measured three times (one reading of each of three wells per culture), and the average of three sterile media readings was used as a reference and subtracted from each culture value. The optical density measurements were used to estimate pyoverdine production per cell (*i.e*. per capita pyoverdine production) using: RFU / OD_600_. The average of the three technical replicates was used in the analysis. We chose to quantify pyoverdine production at the population level (rather than for individual clones) as it represents the most relevant measure of the *Galleria mellonella* inoculum (see *Galleria mellonella virulence assay* below). We repeated this assay with the non-pyoverdine producing strain to confirm it produced negligible quantities of pyoverdine.

### Galleria mellonella virulence assay

To quantify virulence of both the pyoverdine producing and non-producing strains, we used the insect infection model *Galleria mellonella* (44). Briefly, defrosted samples were diluted 10^5^-fold in M9 salt buffer (3g KH_2_PO_4_, 6g Na_2_HPO_4_, 5g NaCl per litre) before 10µL was injected into each of 20 final instar larvae per sample using a 50µL Hamilton syringe (Hamilton, Nevada, USA). Larvae were then incubated at 37°C and mortality checked 18 hours post-injection. Larvae were classed as dead when mechanical stimulation of the head caused no response (45). M9-injected and non-injected controls were used to confirm mortality was not due to injection trauma or background *G. mellonella* mortality; >10% control death was the threshold for re-injecting (no occurrences). Pyoverdine producing and non-producing strains were injected into different *G. mellonella* groups.

### Quantifying Pseudomonas aeruginosa density

The density of *P. aeruginosa* in the common garden environment was quantified by plating onto agar. To do this, samples were defrosted, serially diluted with M9 salt buffer and plated onto KB agar. After 48 hours incubation at 28°C colonies were counted and the number of colony forming units (CFU) standardised to CFUs per mL. Note we only quantified the density of the pyoverdine producing strain.

### Statistical analysis

The effect of copper on per capita pyoverdine production (RFU / OD_600_) and density (CFU/mL) of *P. aeruginosa* populations was tested using linear mixed effects models (LMEM) with copper and time as explanatory variables (both factors), as well as their 2-way interaction. To determine how copper affected virulence we used a binomial generalised linear mixed model (GLMM), with number of *G. mellonella* dead versus alive as binomial response variable, and copper and time as explanatory variables, as well as their two-way interaction. To analyse the combined effect of pyoverdine production and density on virulence, we used a similar framework and included pyoverdine production, total CFU and time, along with a two-way pyoverdine-time interaction, as explanatory variables. In all analyses, pyoverdine production and population density were log_10_-transformed to improve normality, and random intercepts were fitted for individual replicates to account for non-independency of observations over time. For all analyses, we used the ‘*DHARMa’* package (46) to check residual behaviour, after which the most parsimonious model was arrived at by sequentially deleting terms and comparing model fits using *χ* -tests. These were followed by Tukey post hoc comparisons adjusted for multiple comparisons using the Benjamini and Hochberg method. All analyses were carried out in R version 3.3.3 (47), with the ‘*lme4’* package used for the LMEMs and GLMMs (48).

## Results

### Copper selected for increased pyoverdine production

Here we tested the evolutionary consequences of copper stress on *Pseudomonas aeruginosa* by incubating it with copper for either two or twelve days, and then without copper for one day. All results shown here therefore display the evolutionary consequences of copper stress, as the phenotypic effects are removed by growing populations in copper-free common garden environment. Unless otherwise stated, data from the non-pyoverdine producing strain are only shown in the final section of the results. In line with previous findings (3), pyoverdine production evolved to be significantly greater as a function of copper concentration (copper main effect: *χ*^*2*^=35.4, d.f.=2, p<0.001). This effect was consistent across the two- and twelve-day treatments; however, production was significantly greater after twelve days compared to two days (time main effect: *χ*^*2*^=161, d.f.=1, p<0.001; Fig. 2). Pyoverdine production was significantly higher in the high copper treatment compared to both the control (two days: p<0.001; twelve days: p<0.001) and the low copper treatment (two days: p=0.032; twelve days: p=0.032). Likewise, the low copper treatment was always significantly higher than the control (two days: p=0.004; twelve days: p=0.004). The non-independence of observations between day two and day twelve samples accounted for only a small amount of the total variation (SD = 0.029) in this model.

**Figure 2.**
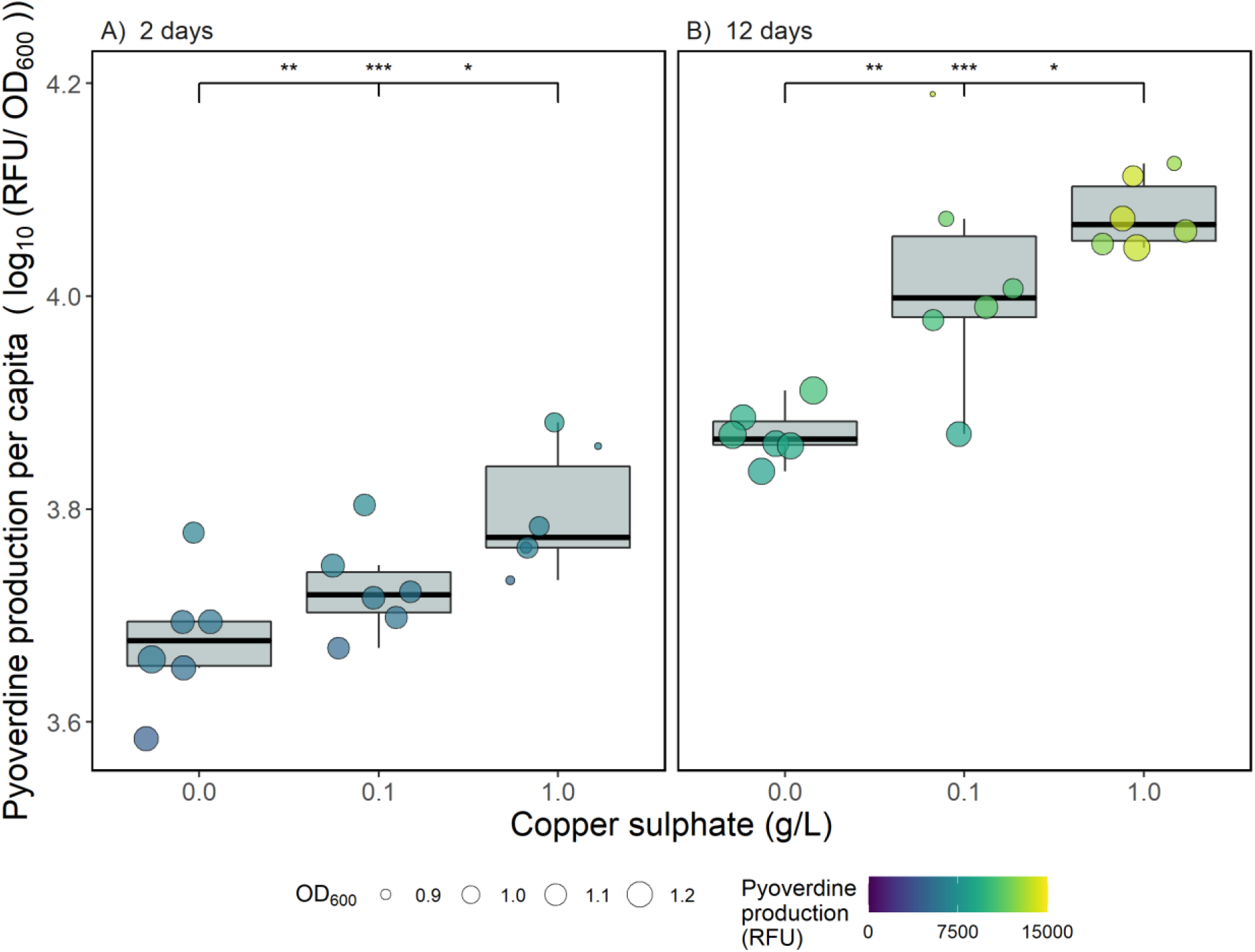
Per capita pyoverdine production (log_10_-transformed standardised fluorescence units per OD_600_) by *Pseudomonas aeruginosa* populations after growth in different concentrations of copper for (A) two days + one day in the absence of copper, or (B) twelve days + one day in the absence of copper. Circles show individual replicates (n = 6), with colour indicating total pyoverdine production (relative fluorescence units) and size showing optical density at 600nm. Asterisks indicate significant differences between groups (*** = 0.001, ** = 0.01, * = 0.05, NS = non-significant), with the left value comparing the control to the low copper treatment, the middle value comparing the control to the high copper treatment and the right value comparing the low and high copper treatments.

### Virulence is higher in populations exposed to copper and this effect increases with time

To test whether copper stress caused differences in *P. aeruginosa* virulence, populations were assayed using the *Galleria mellonella* virulence model. Copper significantly increased virulence (copper main effect: *χ*^*2*^=20.8, d.f.=2, p<0.001) and this effect increased with time (copper-time interaction: *χ*^*2*^=6.90, d.f.=2, p=0.031; Fig. 3). After two days, virulence was significantly higher in the two copper treatments compared to the control (p<0.001 for both comparisons) but these treatments did not differ themselves (p=0.34). After twelve days virulence was again significantly higher in the two copper treatments than the control (p<0.001 for both comparisons), however the high copper treatment was found to be significantly more virulent than the low copper treatment (p=0.027). We note these treatment effects were found despite the random effect of repeatedly measuring the same population accounting for a large amount of the variation (SD = 0.42).

**Figure 3.**
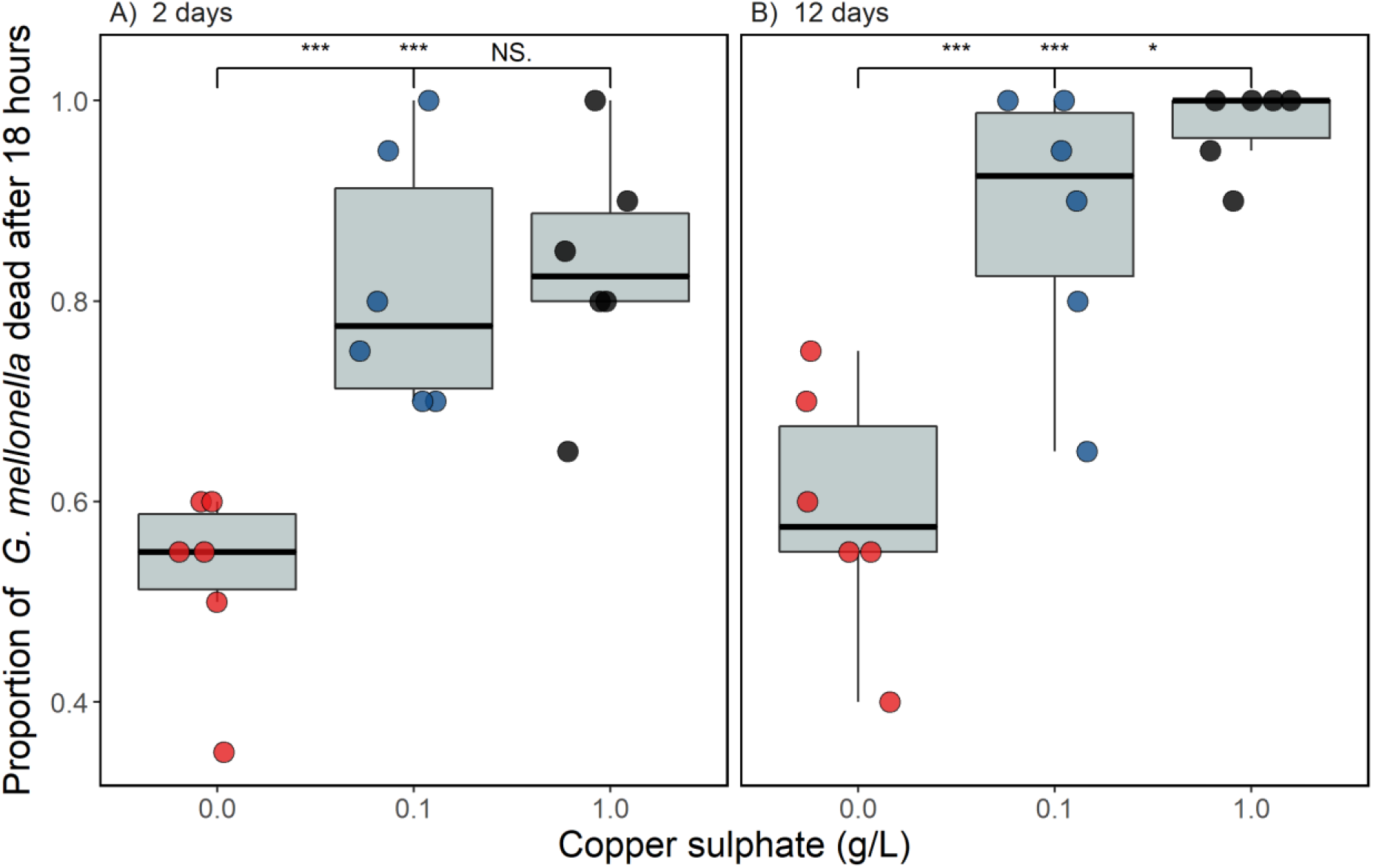
The proportion of *Galleria mellonella* dead 18 hours after being injected with *Pseudomonas aeruginosa* grown in the presence or absence of copper (A) two days + one day in the absence of copper, or (B) twelve days + one day in the absence of copper. 20 *G. mellonella* were injected per replicate (circles show individual replicates; n = 6 per unique treatment combination). Asterisks indicate significant differences between groups (*** = 0.001, ** = 0.01, * = 0.05, NS = non-significant), with the left value comparing the control to the low copper treatment, the middle value comparing the control to the high copper treatment and the right value comparing the low and high copper treatments.

### The effect of copper on population density differs over time

The effect of copper on population density differed as function of time (copper-time interaction: *χ*^*2*^=20.5, d.f.=2, p<0.001; Fig. 4). After two days, population densities did not significantly differ between any copper treatment (p=>0.060 for all contrasts), whereas after twelve days of growth density was significantly lower in the two copper treatments compared to the control (high copper - control: p=0.013 and low copper - control: p=0.002) but did not differ themselves (p=0.48). The effect of measuring the same populations at two time points did not explain any of the overall variation in this model (SD = 0.0), but to account for non-independency of observations it was retained in the model.

**Figure 4.**
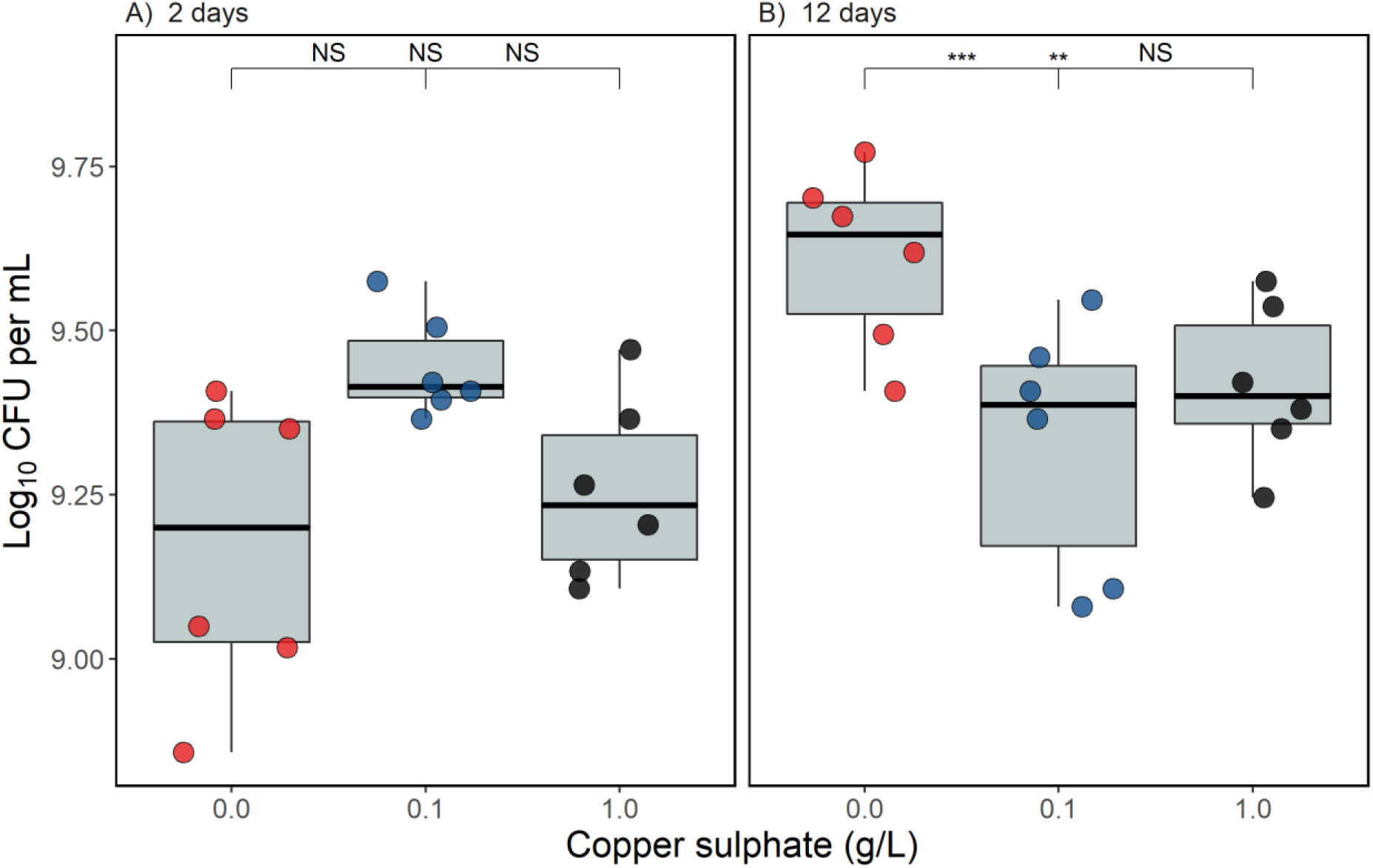
The density (log10 CFU mL^-1^) of *Pseudomonas aeruginosa* populations incubated with copper for (A) two days + one day in the absence of copper, or (B) twelve days + one day in the absence of copper. Asterisks indicate significant differences between groups (*** = 0.001, ** = 0.01, * = 0.05, NS = non-significant), with the left value comparing the control to the low copper treatment, the middle value comparing the control to the high copper treatment and the right value comparing the low and high copper treatments.

### Virulence is associated with increased pyoverdine production rather than pathogen load

Next, we tested whether increased virulence was associated with increased pyoverdine production by replacing ‘copper’ as an explanatory variable in our GLMM with *per capita* pyoverdine production and included density as a covariate in this model. Despite there being substantial variation across populations in virulence (SD = 0.86), virulence increased as a function of pyoverdine production (pyoverdine main effect: *χ*^*2*^=13.9, d.f.=1, p<0.001; Fig. 5), and this effect did not change with time (pyoverdine-time interaction: *X*^*2*^=0.009, d.f.=1, p=0.93) although virulence was higher after twelve days (time main effect: *X*^*2*^=5.40, d.f.=1, p=0.02). Population density had no significant effect on virulence (main effect of density in GLMM: *X*^*2*^=1.09, d.f.=2, p=0.30).

**Figure 5.**
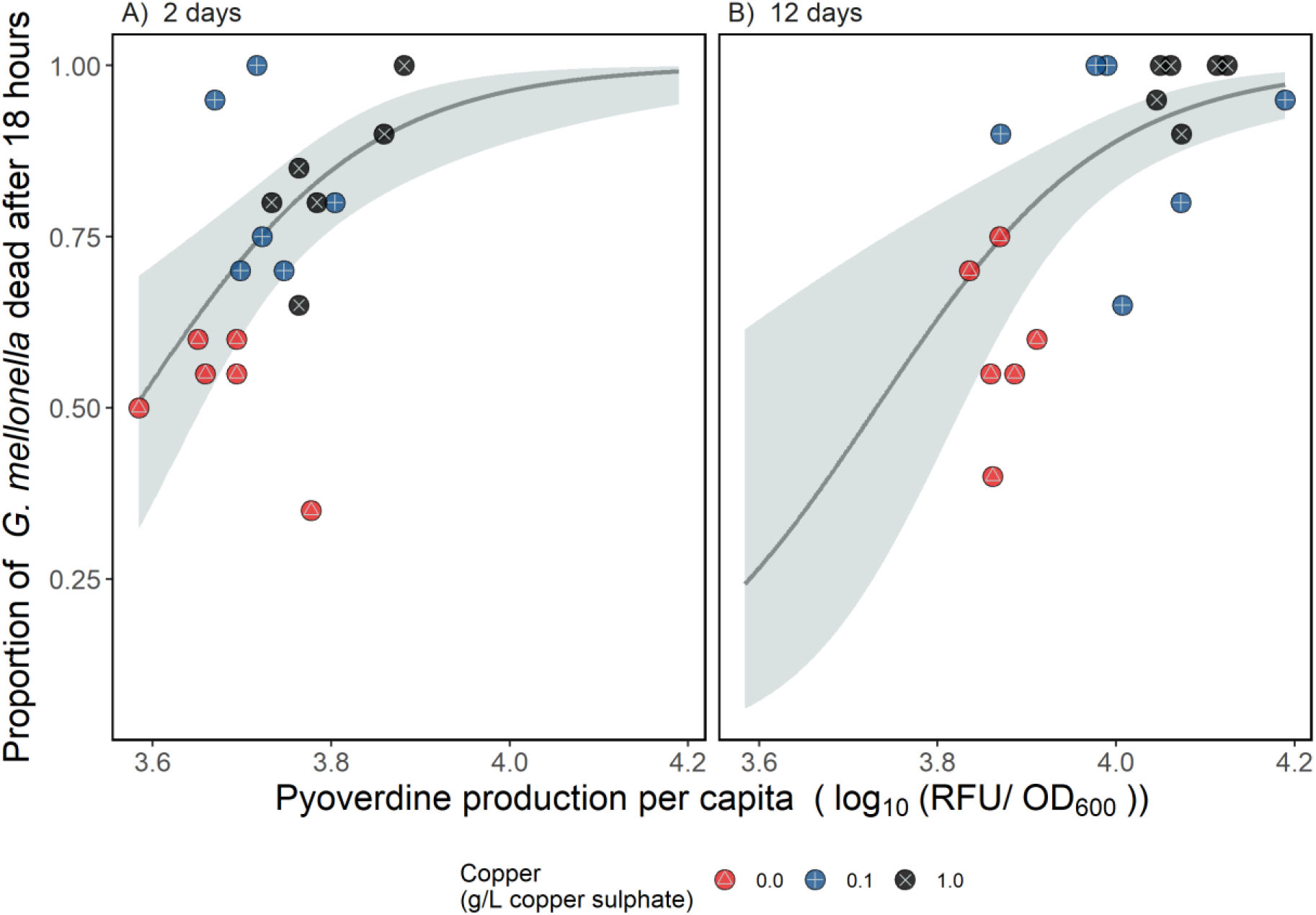
The relationship between per capita pyoverdine production (log_10_-transformed relative fluorescence units per OD_600_) and virulence of *P. aeruginosa* populations grown with copper for (A) two days + one day in the absence of copper, or (B) twelve days + one day in the absence of copper. Virulence was quantified using the *Galleria mellonella* infection model and expressed as the proportion of *G. mellonella* dead (out of 20) 18 hours after injection. Individual replicates are represented by circles (n = six per treatment); red points containing a Δ represent the control (no copper) treatments, blue points containing a + the low copper treatment, and black points containing a × the high copper treatment. The line shows the best model fit, and the shaded area shows the 95% confidence interval.

### The high copper treatment decreased the virulence of the non-pyoverdine producing strain

Finally, to confirm at least part of the change in virulence was due to pyoverdine production, we tested whether a non-pyoverdine producing mutant strain of *P. aeruginosa* evolved under the same copper regime exhibited similar changes in virulence. The fluorescence assay used to detect pyoverdine suggested that the mutant strain still produced some pyoverdine (mean = 431 ± 91.6 SD RFU), but this was on average 94.5% lower than populations of the wild-type producer (mean = 7763 ± 2979 SD RFU), and therefore deemed negligible (data not shown). Once again, we found a significant effect of copper on virulence (copper main effect: *χ*^*2*^=66.6, d.f.=2, p<0.001; Fig. 6) however, this time the effect was negative, and decreased with time (copper-time interaction: *χ*^*2*^=6.45, d.f.=2, p=0.040; Fig. 6). Despite a slight decrease in the effect of copper over time, we found the same qualitative pattern in the virulence of the non-pyoverdine producing strain across copper treatments after two and twelve days: virulence was not significantly different between the low copper treatment and the control (two days: p=0.087; twelve days: p=0.48), but was significantly lower in the high copper treatment compared to both the low copper treatment and the control (p=<0.001 for all contrasts; Fig. 6). As with the virulence model for the pyoverdine producing strain, the random effect of repeatedly measuring the same population accounted for a large amount of the variation (SD = 0.44). Note that injections with pyoverdine producing and non-pyoverdine producing strains were done in different batches of *G. mellonella*, and so statistical comparison between the results from the wild-type and mutant infections is inappropriate.

**Figure 6.**
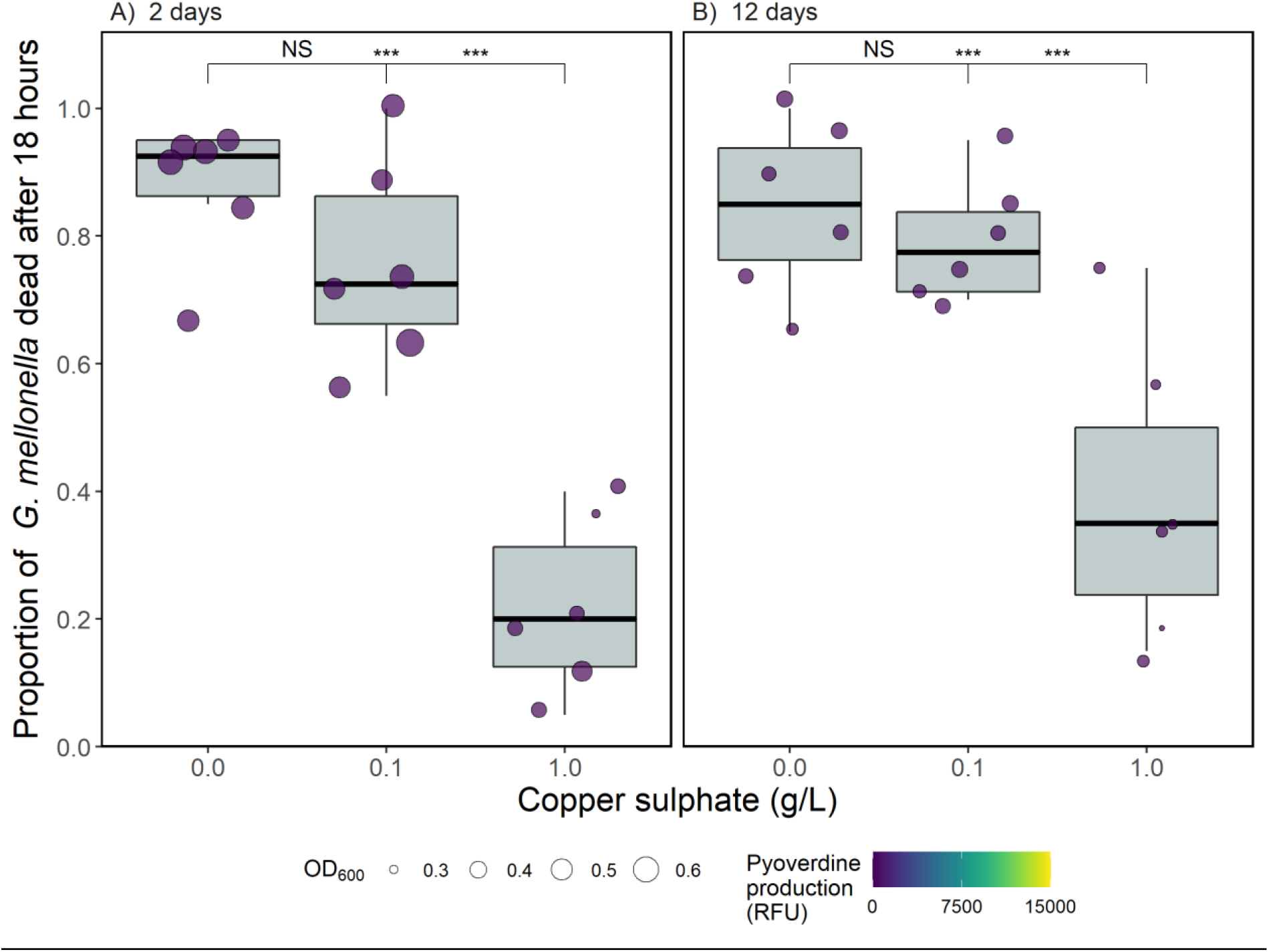
The proportion of *Galleria mellonella* dead 18 hours after being injected with an isogenic non-pyoverdine producing strain of *P. aeruginosa*grown in the presence or absence of copper for (A) two days (+ one day in the absence of copper) or (B) twelve days (+ one day in the absence of copper). 20 *G. mellonella* were injected per replicate. Circles show individual replicates (n = 6), with size showing optical density at 600nm and colour indicating total pyoverdine production (standardised fluorescence units) to demonstrate the mutant did indeed not produce any pyoverdine. Asterisks indicate significant differences between groups (*** = 0.001, ** = 0.01, * = 0.05, NS = non-significant), with the left value comparing the control to the low copper treatment, the middle value comparing the control to the high copper treatment and the right value comparing the low and high copper treatments.

## Discussion

Here, we experimentally tested whether copper causes the evolution of increased siderophore-mediated virulence in *Pseudomonas aeruginosa*. We found copper to select for increased per capita pyoverdine production, which was associated with greater levels of death in the *Galleria mellonella* infection assay. Moreover, pyoverdine production was found to increase with time, such that populations exposed to copper for twelve days were more virulent than those exposed for two days. In addition to this overall increase in virulence with time, high copper caused significantly greater virulence than low copper in the twelve-day treatment but not in the two-day treatment. As a result, the twelve-day high copper treatment demonstrated the greatest virulence. Furthermore, we found that when using a non-pyoverdine producing mutant strain of *P. aeruginosa*, high copper reduced virulence, and low copper had little effect on virulence. Our results therefore demonstrate that copper stress increases pyoverdine production and consequently virulence, and that this effect increases with exposure time and copper concentration.

Finding copper-mediated increases in pyoverdine production is consistent with a role of pyoverdine in detoxification of copper in *P. aeruginosa* populations (49). Here, we show that a prolonged need for copper detoxification leads to the continued evolution of greater siderophore production. The resulting cost of these genotypic changes to a metabolically costly trait is a likely explanation for the reduced densities in the twelve-day copper treatments. We note that our finding of increased pyoverdine production with copper differs to that of previous work (8). Likely explanations for this are the fewer observed generations in our study (40 opposed to 200), culturing under static (versus shaken) conditions, which favours public goods production, and transfers occurring every two days compared to daily. In addition to copper, pyoverdine production has been shown to be up-regulated in response to Al^3+^, Ga^3+^, Mn^2+^ and Ni^2+^ (49). This suggests that these metals may also increase *P. aeruginosa* virulence, and may cause selection for increased production over longer exposure periods. Furthermore, it is likely that mixtures of metals, which are frequently encountered in polluted environments, might additively select for pyoverdine production and thereby virulence.

The positive association between pyoverdine production and virulence is consistent with previous findings, including in murine models (50). As well as directly aiding pathogen growth within a host by increasing iron uptake, pyoverdine production can increase virulence by causing the upregulation of additional virulence factors such as Exotoxin A and PrpL protease (51, 52). Finding per capita pyoverdine production to be a better predictor of virulence than population density in this system shows the cost of evolving greater production on density is inconsequential in a host. Furthermore, finding the virulence of the non-pyoverdine producing mutant to not be affected by low copper, and to be decreased by high copper, strongly suggests that virulence has changed as a consequence of the evolutionary change in pyoverdine production. However, we note that in addition to pyoverdine production, copper stress has been shown to change the expression of over 300 genes in *P. aeruginosa* (53), and it is plausible these also have implications for virulence.

In addition to *P. aeruginosa*, toxic metals have also been shown to induce the production of (non-pyoverdine) siderophores in other species (10), with siderophore production being shown to increase proportionally with toxic metal pollution in natural microbial communities (9). As the copper concentrations used in this study were chosen for their environmental relevance, we suggest our findings could be relevant to natural communities inhabiting polluted environments. We note that remediation techniques, principally lime addition, are used in some metal-polluted areas to reduce their toxic effect (54), and this can lower community siderophore production (55). Extending experiments such as those described here to the level of natural communities could shed light on the consequences of metal pollution and metal remediation on bacterial siderophore production and virulence.

In conclusion, we experimentally show that copper can increase the virulence of the pathogen *Pseudomonas aeruginosa* by selecting for increased production of a metal-detoxifying siderophore. We therefore demonstrate a direct link between toxic metal stress and virulence in an opportunistic pathogen of significant clinical importance. Furthermore, we show that the effect of metal exposure on virulence increases with exposure time and copper concentration. This raises further concern for the effect of ever-increasing metal pollution on bacterial pathogens, and highlights further work is needed to understand the role of metals in bacterial virulence.

## Data accessibility

All data and code will be made publically available upon acceptance. Currently, data can be found on Mendeley Data at https://data.mendeley.com/v1/datasets/yypr3yfr86/draft?preview=1, and the R code to reproduce all analysis and create all data figures at https://data.mendeley.com/v1/datasets/hbf9ht7d8h/draft?preview=1.

## Author contributions

LL conceived the study, carried out experimental work, statistical analysis of the results and drafted the manuscript. EH helped with the analysis of the results. All authors helped with experimental design and critically revised the manuscript.

## Acknowledgments

LL would like to thank the NERC FRESH GW4 award no. NE/R011524/1, EH the UKRI\ Future Leaders Fellowship award MR/V022482/1, AB the NERC award NE/V012347/1, and MV the NERC award NE/T008083/1.

## Notes

### Competing Interest Statement

The authors have declared no competing interest.

